# Mathematical modeling of the molecular switch of TNFR1-mediated signaling pathways using Petri nets

**DOI:** 10.1101/2021.11.02.466901

**Authors:** Leonie K. Amstein, Jörg Ackermann, Jennifer Hannig, Ivan Ðikić, Simone Fulda, Ina Koch

## Abstract

The paper describes a mathematical model of the molecular switch of cell survival, apoptosis, and necroptosis in cellular signaling pathways initiated by tumor necrosis factor 1. Based on experimental findings in the current literature, we constructed a Petri net model in terms of detailed molecular reactions for the molecular players, protein complexes, post-translational modifications, and cross talk. The model comprises 118 biochemical entities, 130 reactions, and 299 connecting edges. Applying Petri net analysis techniques, we found 279 pathways describing complete signal flows from receptor activation to cellular response, representing the combinatorial diversity of functional pathways.120 pathways steered the cell to survival, whereas 58 and 35 pathways led to apoptosis and necroptosis, respectively. For 65 pathways, the triggered response was not deterministic, leading to multiple possible outcomes. Based on the Petri net, we investigated the detailed *in silico* knockout behavior and identified important checkpoints of the TNFR1 signaling pathway in terms of ubiquitination within complex I and the gene expression dependent on NF-κB, which controls the caspase activity in complex II and apoptosis induction.

## Introduction

The tumor necrosis factor receptor 1 (TNFR1) controls pivotal cellular processes involved in immunity and developmental processes (Walczak & Kantari, 2011). TNFR1 mediates signaling pathways, which induce opposing cellular responses from initiation of gene expression to two forms of cell death, apoptosis and necroptosis (Walczak, 2011; Pasparakis & Vandenabeele, 2015). Apoptosis has long been viewed as the only form of cell death, which is initiated by the cell itself. Whereas apoptosis is a well-known and well-studied pathway, the regulation and function of the necroptosis pathway has just recently been discovered and is still under study (Dhuriya & Sharma, 2018; Degterev, 2005). Necroptosis describes a cell death mode that exhibits the phenotype of necrosis, although it is ordered and controlled like apoptosis (Vandenabeele *et al*, 2010). Alike necrosis, necroptosis features a form of cellular explosion, releasing the cellular content into the cell surrounding and initiating inflammation in the tissue (Vandenabeele *et al*, 2010). On the contrary, cells that undergo apoptosis recycle most of the cellular molecules to reserve the energy and slowly digest themselves without inducing an inflammatory response in the surrounding cells (Reed & Green, 2011). It has been reported that necroptosis seems to play a crucial role in nonalcoholic fatty liver disease, nonalcoholic steatohepatisis, and liver cancer (Schwabe & Luedde, 2018).

Alternatively to cell death, the activation of nuclear factor κ-light-chain-enhancer of activated B cells (NF-κB) initiates the gene expression of mainly pro-inflammatory and anti-apoptotic operating genes (Pasparakis & Vandenabeele, 2015). Thus, the NF-κB pathway is often referred to as the survival pathway triggered by TNFR1 stimulation (Walczak & Kantari, 2011). A permanent activation of NF-κB can result in chronical inflammation and promote the formation of tumors (DiDonato *et al*, 2012). In cancer cells, the gene expression is often permanently active, for example, by a disruption of the TNFR1 signaling pathway, such that the cells exhibit a resistance against cell death induction. Anticancer therapy aims to induce cell death in cancer cells often by triggering apoptosis pathways (Fulda *et al*, 2010; Fulda, 2011; Fulda & Vukic, 2012, Fulda, 2013) and therapeutic exploitation of necroptosis (Fulda, 2014).

The regulation of the opposing signaling cascades is often considered as the molecular switch. Receptor-interacting protein 1 (RIP1) seems to have a pivotal function in modulating the controversial outcomes since it is an essential signaling node in all pathways, see Fig 1. The activity and function of RIP1 is sensitively controlled (Pelzer *et al*, 2006), for example, by post-translational modifications, such as phosphorylation and ubiquitination. During ubiquitination, ubiquitin (Ub) covalently attaches Ub molecules to substrate proteins, forming chains of different linkage types (Ikeda & Dikic, 2008) and assigning specific functions to the respective proteins (Grabbe *et al*, 2011). Linear Ub chains influence the modulation and control of activity in signal transduction (Walczak *et al*, 2012; Kensche *et al*, 2012, Declercq et al, 2009). The Ub system may have a promising therapeutic potential similar to the post-translational modification of phosphorylation mediated by kinases (Hoeller & Dikic, 2009; Fulda *et al*, 2012).

**Figure 1:**
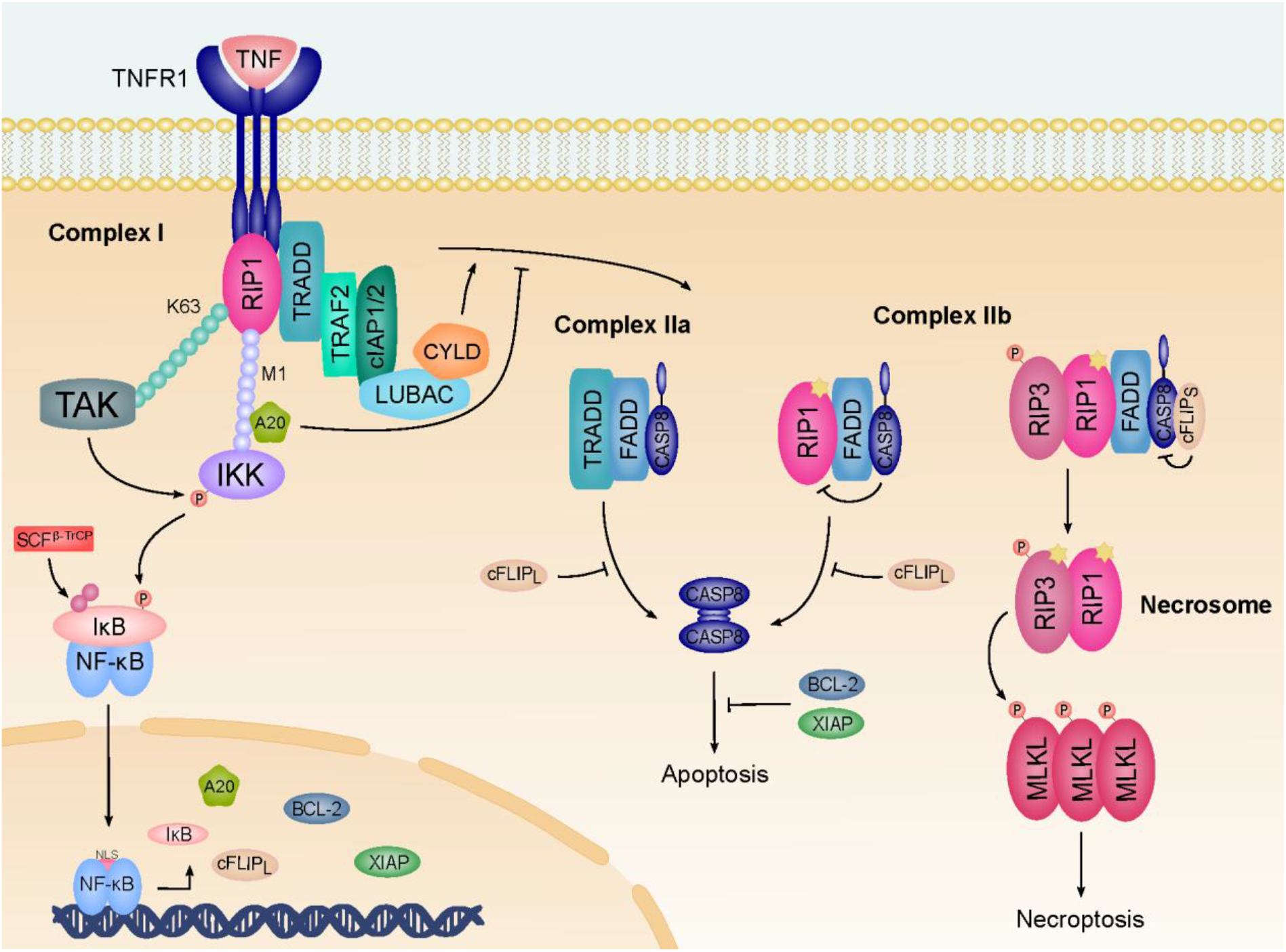
The TNFR1 signal transduction pathway. Upon engagement of TNFR1, complex I is rapidly formed and mediates the signaling to NF-κB activation. The ubiquitination mediated by E3 ligases, like cellular inhibitor of apoptosis protein 1 (cIAP1) or cellular inhibitor of apoptosis protein 2 (cIAP2) and linear ubiquitin chain assembly complex (LUBAC), promotes the association of complex I. The Ub modification is required for full activation of the inhibitor of NF-κB (IκB) and subsequent NF-κB activation. Activated NF-κB in the nucleus initiates the expression of target genes like IκB, A20, cellular FLICE-inhibitory protein (cFLIP_L_), B-cell lymphoma 2 (BCL-2), and X-linked inhibitor of apoptosis protein (XIAP). A20 is a deubiquitinating enzyme (DUB), which is reported to cleave lysine 63 (K63) chains while protecting methionine 1 (M1) chains from cleavage. The deubiquitination by CYLD destabilizes the complex and promotes the formation of complex II in the cytosol. Complex IIa associates caspase 8 (CASP8), while complex IIb additionally binds RIP1. cFLIP_L_ reduces, but does not fully inhibit, caspase activity, which leads to RIP1 and RIP3 cleavage and inhibits apoptosis and necroptosis. cFLIP_S_ fully inhibits caspase activity and promotes the formation of the necrosome. Autophosphorylation of RIP3 allows the recruitment and phosphorylation of MLKL, which subsequently forms active oligomers and translocates to the plasma membrane to induce necroptosis.

Although high-throughput technologies have provided many experimental data, there is a lack regarding the quality, quantity, and completeness of the data. Applying computational systems biology can provide informationon on the system-wide behavior without knowing kinetic parameters. Computational models are powerful approaches to represent and understand the complexity of biological systems and systematically analyze them. These analyses gain new insights of regulation, reveal implications in diseases and pathologies, and give useful implications for potential targets for therapeutic treatment (Kitano, 2002, 2004). The emerging experimental procedures, in coordination with improved computational methods, seem promising for the analysis of signaling pathways also with regard to therapeutic intervention and drug treatment (Saez-Rodriguez *et al*, 2015). The data available and the questions addressed determine the modeling approach to be applied. These approaches cover kinetic or stochastic quantitative models, for example, systems of ordinary differential equations (ODEs) (e.g., Heinrich & Rapoport, 1973), qualitative models as Boolean models (Aldridge *et al*, 2006; Wang *et al*, 2012), or semi-quantitative models, such as Petri nets (PNs) (Reisig, 1985; Murata, 1989). PNs allow for qualitative discrete modeling as well as for quantitative, continuous modeling. They have been widely applied to model biological pathways at different scales of abstraction (Koch *et al*, 2005; Formanowicz *et al*, 2006; Sackmann *et al*, 2007; Koch *et al*, 2011; Minervini *et al*, 2014; Koch *et al*, 2017; Jacobsen *et al*, 2020). Additionally, PNs provide a simplified and clear user-friendly visualization of the model graph (e.g., Einloft *et al*, 2013; Balazki *et al*, 2015).

The TNFR1 signaling pathway has often been a subject of mathematical modeling (Mitchell *et al*, 2016), thus, revealing dynamics, regulations, and crosstalk to other pathways of the NF-κB pathway (Basak *et al*, 2012; Cheng *et al*, 2012). On the one hand, the NF-κB regulation is well characterized and has often been a subject of quantitative modeling approaches, such as an ODE-based model of the NF-κB signaling module (Hoffmann *et al*, 2002). According to new measured values and estimated parameters, there exist various adaptations and further developments of this model (Lipniacki *et al*, 2004; Kearns *et al*, 2006; Rangamani & Sirovich, 2007; Tay *et al*, 2010; Sheppard *et al*, 2011; Mothes *et al*, 2015). On the other hand, new insights often supersede older views of the pathway regulation and initiate the development of, for example, a hybrid PN of NF-κB activation and regulation of gene expression (Peng *et al*, 2010). A Boolean model describes the interplay between NF-κB activation, apoptosis, and necroptosis, following the stimulation of TNFR1 and FAS receptor (Calzone *et al*, 2010). Schlatter *et al*. proposed a Boolean model of the processes of apoptosis, which considers several stimuli (Schlatter *et al*, 2009). Schliemann *et al*. have merged two existing models to an ODE-based model with pro- and anti-apoptotic responses of TNFR1 signaling (Schliemann *et al*, 2011). Melas *et al*. introduced a hybrid model, covering the stimulation of seven receptors and 22 cytokine stimuli in immunological pathways (Melas *et al*, 2011). All these models focus on specific processes or stimuli, not considering the entire molecular switch between cell survival, apoptosis, and necroptosis.

In this paper, we are interested in an exhaustive modeling of the molecular switch behavior of the TFNR1-induced signaling pathway, covering the NF-κB pathway, apoptosis, and necroptotic processes. Here, we developed a semi-quantitative PN model and applied invariant-based methods and *in silico* knockout analysis to investigate and discuss the system’s behavior of the PN. This includes a detailed discussion of the molecular switch behavior.

## Results

### The Petri net model of signaling processes of cell survival, apoptosis, and necroptosis

In the following, we refer to the PN terminology, which we explain in detail in Section “*Materials and Methods*”. **Fig 1** schematically illustrates the molecular processes of TNFR1 signal transduction commenced by the stimulation of the TNF receptor and followed by the formation of complex I and a diversity of consecutive and concurrent molecular processes. An example is the translocation of NF-κB into the nucleus, which facilitates gene expression activity and transcription of proteins like IκB, A20, cellular FLICE-inhibitory protein (cFLIP_L_), B-cell lymphoma 2 (BCL-2), and X-linked inhibitor of apoptosis protein (XIAP). The transcription of these proteins affects, e.g., the regulation of the TNFR1 signaling pathway. The formation of complex IIa, complex IIb, and the necrosome may induce either apoptosis or necroptosis.

Based on the prcesses illustrated in **Fig 1**, we constructed a PN model to analyze the broad combinatorial spectra of signaling pathways. The model comprises stoichiometry relations for well-studied processes in combination with the abstraction of a simple transition for processes with unknown stoichiometry or controversial experimental findings. **Fig 2** represents the PN model with 118 places, 130 transitions, and 299 edges. For the list of transitions, places, and label abbreviations, we refer to **Table S1, Table S2**, and **Table S3**, respectively (Appendix). Signaling cascades towards NF-κB activation, apoptosis, and necroptosis are highlighted blue, green, and red, respectively. A dot in a circle indicated a place with one token in the initial marking. Gray circles represent a place that appears at several locations of the network layout. On left side, the layout separately shows the synthesis of 26 housekeeping proteins that are required for maintening the basic cellular function. The input and output transitions are labeled according to their biological meaning. All other transitions are consecutively numbered. All input transitions represent syntheses of proteins.The output transitions model the diverse cellular outcomes, like apoptosis, necroptosis, or survival as well as degradation and dissociation processes for proteins and protein complexes, respectively. The places were labeled according to the biological meaning, e.g., a protein, a modified protein, or a protein complex.

**Figure 2:**
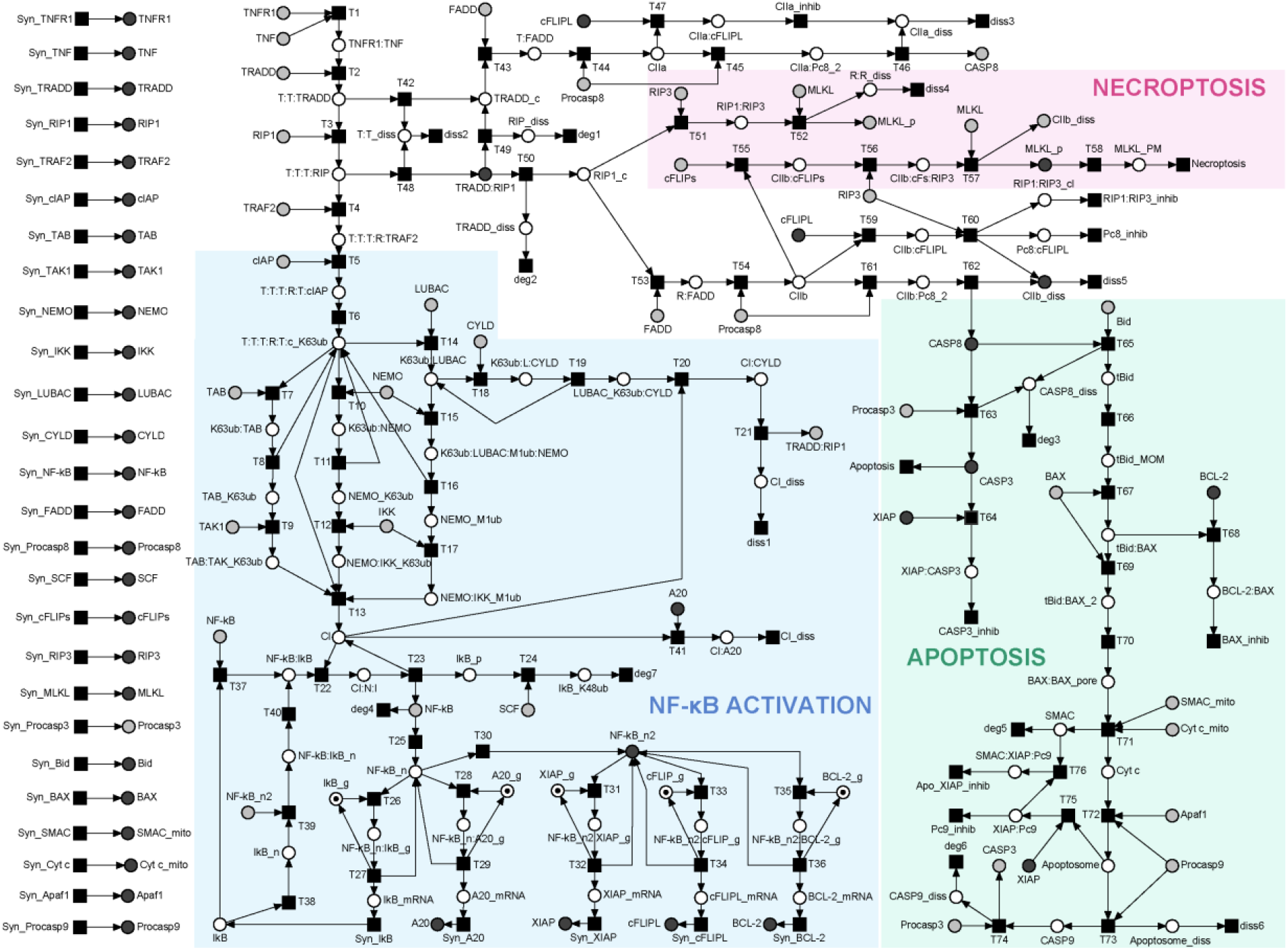
The PN model of TNFR1 signal transduction. The PN consists of 118 places, 130 transitions, and 299 edges. The essential processes of NF-κB activation, apoptosis, and necroptosis are highlighted blue, green, and red, respectively. Logical places are depicted in gray. The initial marking is represented by one token assigned to places *IκB_g, A20_g, XIAP_g, cFLIP_g*, and *BCL-2_g* (*g* stands for gene) for each PI.

To ensure correctness and completeness of the model to the greatest possible extent, we apply the invariant analysis.

### Place invariants reflect substance conservation

The five place invariants (PIs) of the PN represent the conservation of the proteins IκB, A20, XIAP, cFLIPL, and BCL-2, all containing two places.

### Transition invariants reflect basic dynamic patterns

The PN is covered by 48 transitions invariants (TIs). For the list of TIs and their biological interpretations, we refer to **Table S4** (Appendix). Exemplarily, **Fig 3** highlights TI_2_, which describes a signal flow from TNFR1 activation to apoptosis. It comprises the formation of complex I, dissociation via CYLD, formation of complex IIb, and extrinsic activation of caspase 3. TIs are functional submodules but not each TI represents a complete signaling pathway, for example, TI_15_ and TI_9_, see **Fig S1** and **Fig S2**, respectively (Appendix). 33 TIs of the total number of 48 TIs are such incomplete signaling pathways (see Table **S4** in the Appendix, bold-faced TI numbers). The remaining 15 TIs are Manatee invariants (MIs), describing complete signaling pathways from the receptor activation to the cell response (Amstein *et al*, 2017), see Section “*Materials and Methods*”.

**Figure 3:**
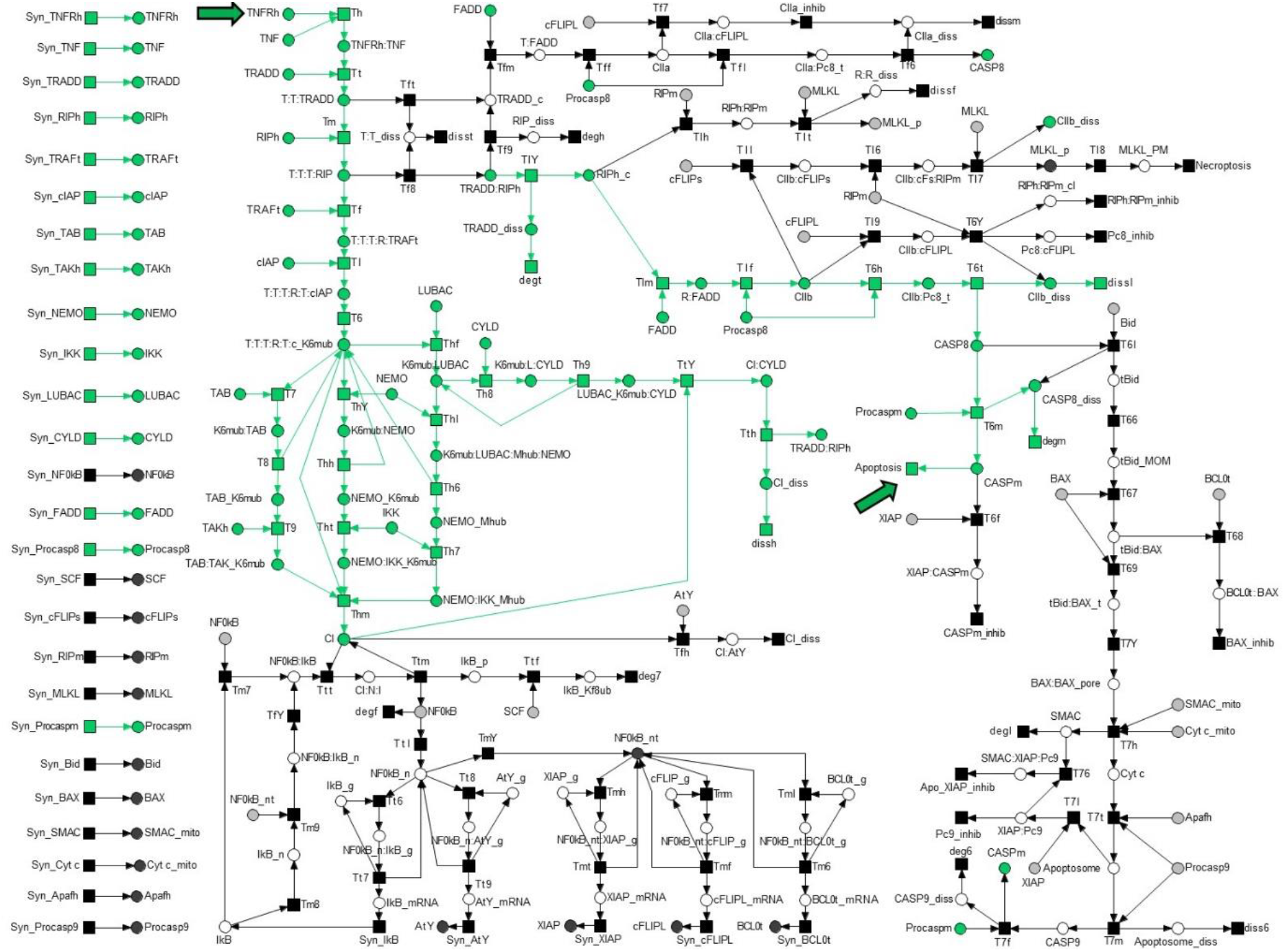
The TI_2_-induced subnetwork. The TI_2_-induced subnetwork is highlighted green in den PN model of **Fig 2**. It covers the formation of complex IIb and induction of apoptosis via the activation of CASP3 in the extrinsic pathway. The start and end transition are indicated by green arrows.

### Manatee invariants describe complete signaling pathways from receptor activation to cell response

We found overall 279 MIs (Table **S5** in the Appendix) by linear combination of TIs. Each of the 279 MIs represents a unique pathway of the molecular switch between cell survival, apoptosis, and necroptosis. Exemplarily, **Fig 4** highlights MI_7_ that combines three TIs, TI_9_, TI_15_, and TI_18_. The red signal flow described by TI_15_ requires NF-κB, i.e., a token on place *NF-κB*, as well as a token on place *CI* (complex I). NF-κB is provided by transition *Syn_NF-κB* of the green TI_18_. Complex I is provided by T_13_ of the blue TI_9_. Vice versa, the signal flow described by the blue TI_9_ cannot work without NF-κB in the nucleus, i.e., a token on place *NF-κB_n*. Translocation of NF-κB into the nucleus requires an active transition T_25_ of the red TI_15_. MI_7_ demonstrates typical mutual dependencies of TIs that make isolated TIs nonfunctional.

**Figure 4:**
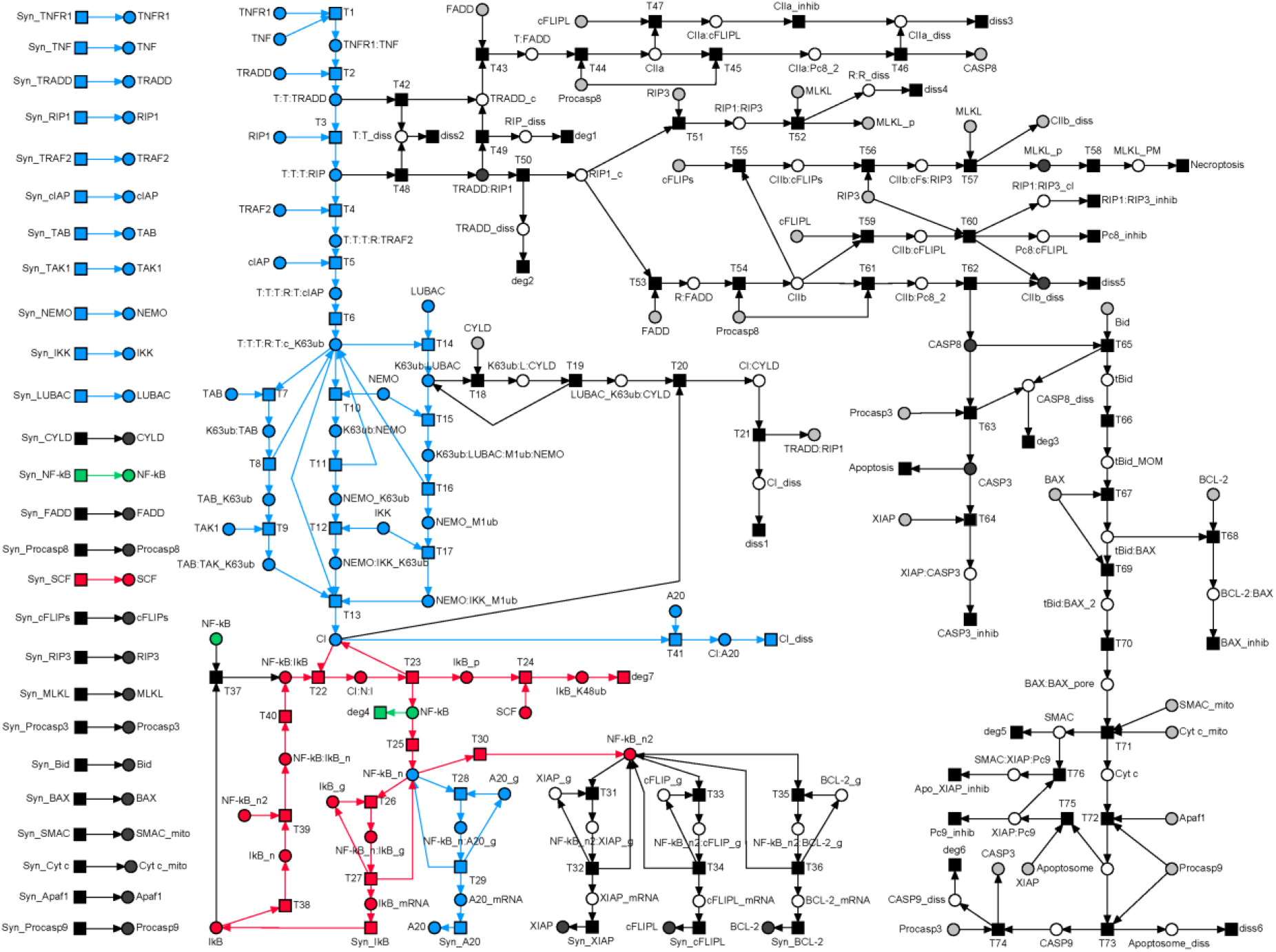
The MI_7_-induced subnetwork of the PN model in Fig 2. The MI_7_-induced subnetwork consists of TI_9_, TI_15_, and TI_18_ highlighted blue, red, and green, repectively.

TI_9_ (**Fig S2**, Appendix) represents the dissected pathway of the A20 feedback regulation in complex I. MI_7_ (**Fig 4**), including TI_9_, determines a complete signal flow, including the A20 feedback loop and covering the signal flow of complex I formation and activation of NF-κB with its translocation into the nucleus and gene expression of IκB and A20. The inhibitor, IκB, terminates gene expression and restores the inhibitory complex of NF-κB and IκB in the cytosol. A20 binds to complex I, leads to the dissociation of the complex, and prevents the formation of complex II. For additional MIs, containing TI_9_ too, see **Fig S3** and **Fig S4** (Appendix).

### Classification of MI-defined signaling pathways

Each of the 279 MIs denotes a complete and unique signaling pathway, see **Table S5** (Appendix). For space reasons, we abstain from a discussion of each individual pathway. We classified the MIs according to their biological outcome, considering the 166 MI-induced subnetworks with a clear classification into a survival, apoptosis, or necroptosis pathway. An ambiguous pathway covers, e.g., the inhibition of MOMP induction, which would result in cell survival and apoptosis induction via the extrinsic pathway. In this special case, MOMP induction is part of the intrinsic pathway, but extrinsic apoptosis induction is still possible. Thus, the MOMP induction would be classified as an apoptosis pathway. This held for 48 MIs of the 113 MIs, so they were all considered for the classification, overall 214 (48 + 166 MIs).

A central property of a pathway is the induced cellular response. The largest fraction of 120 pathways steered the cell to survival, whereas 58 and 35 pathways led to apoptosis and necroptosis, respectively. 65 pathways were neglected because they either could trigger both types of cell death, apoptosis and necroptosis, or represent housekeeping pathways without induction of a specific cellular response. A simple example of a housekeeping pathway is the synthesis and degradation of NF-κB described by TI_18_ highlighted green in **Fig 4**. Note that, TI_18_ also corresponds to MI_18_. For pathways that can trigger both types of cell death, accurate quantitative simulations would be required to determine the stochastic chance of the cell to end up either in apoptosis or necroptosis.

### *In silico* knockouts

The knowledge of the combinatorial diversity of pathways enabled us to estimate the vulnerability of the system to pertubations, caused, for example, by knockouts of proteins. We applied *in silico* knockout analysis to get the number of blocked molecular species downstream of the pathways (Hannig *et al*. 2019). **Fig 5** shows a bar plot of the percentage of the network that becomes inoperable, if we would knockout the synthesis of a specific protein. We ranked the proteins according to the percentage of blocked species. TNF-α (called TNF in **Fig 5**) and TNFR were top-ranked as they are essential upstream in each pathway. Only housekeeping pathways remained unaffected. Components of complex I, e.g., TRADD and RIP, were among high-ranked proteins, too. Proteins of the intrinsic apoptotic branch, necroptosis, and proteins upregulated by NF-κB had more specific functions in the molecular switch and got a lower ranking.

**Figure 5:**
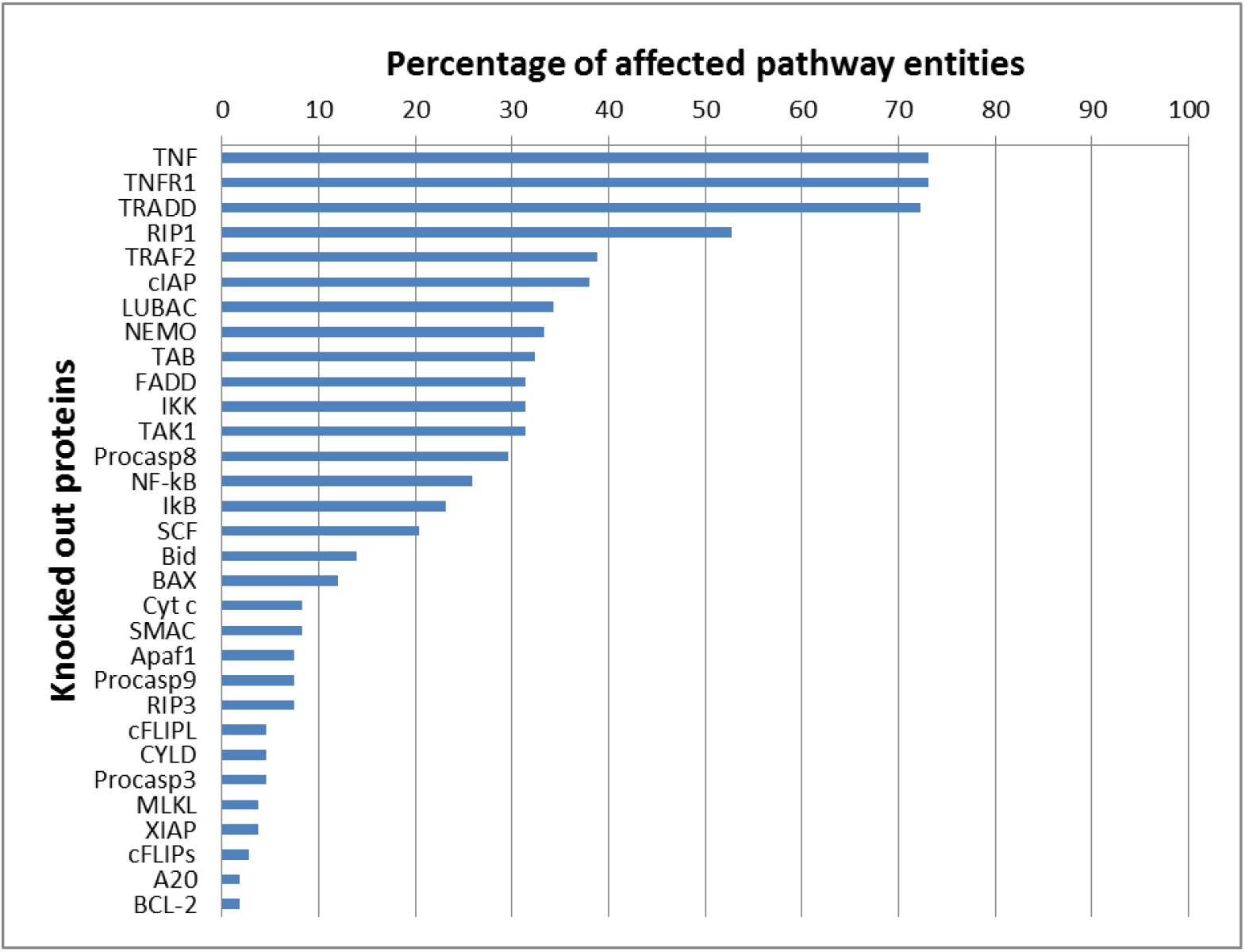
Ranking of the proteins of the TNFR1 signaling pathway. The influence on all other network components was determined based on the *in silico* knockout matrix in **Fig S5** (Appendix). The bar chart displays the relative abundance of affected species for the knockout of all proteins.

### The hierarchical cluster tree

The cluster tree in **Fig 6** outlines the hierarchical organization of function in the signaling pathway. Each leaf of the tree is a protein. To cluster the proteins, we represented a protein by the downstream effect of its knockout, i.e., the set of blocked species. For the detailed knockout matrix, we refer to **Fig S5** (Appendix). Specific branch points were labeled by the characteristic, regulative function of the group of proteins, e.g., ubiquitination in complex I, activation of CASP8, and activation of NF-κB. Three main groups emerged, one for each of the functions apoptosis (green branch), regulation of complex I (blue/purple branch), and necrosis (red branch). Due to crosstalk and feedback, the regulation of complex I was more strongly coupled to apoptosis than to necrosis, leading to a unification of the two branches *Regulation of complex I* and *Apoptosis*. The three proteins of the necroptosis branch, RIP3, MLKL, and cFLIPs, were grouped together very late. Large clusters for the activation of NF-κB or the intrinsic pathway of apoptosis were already complete before RIP3 and MLKL clustered together. The necroptosis branch remained separated from all other functions until the very last clustering step.

**Figure 6:**
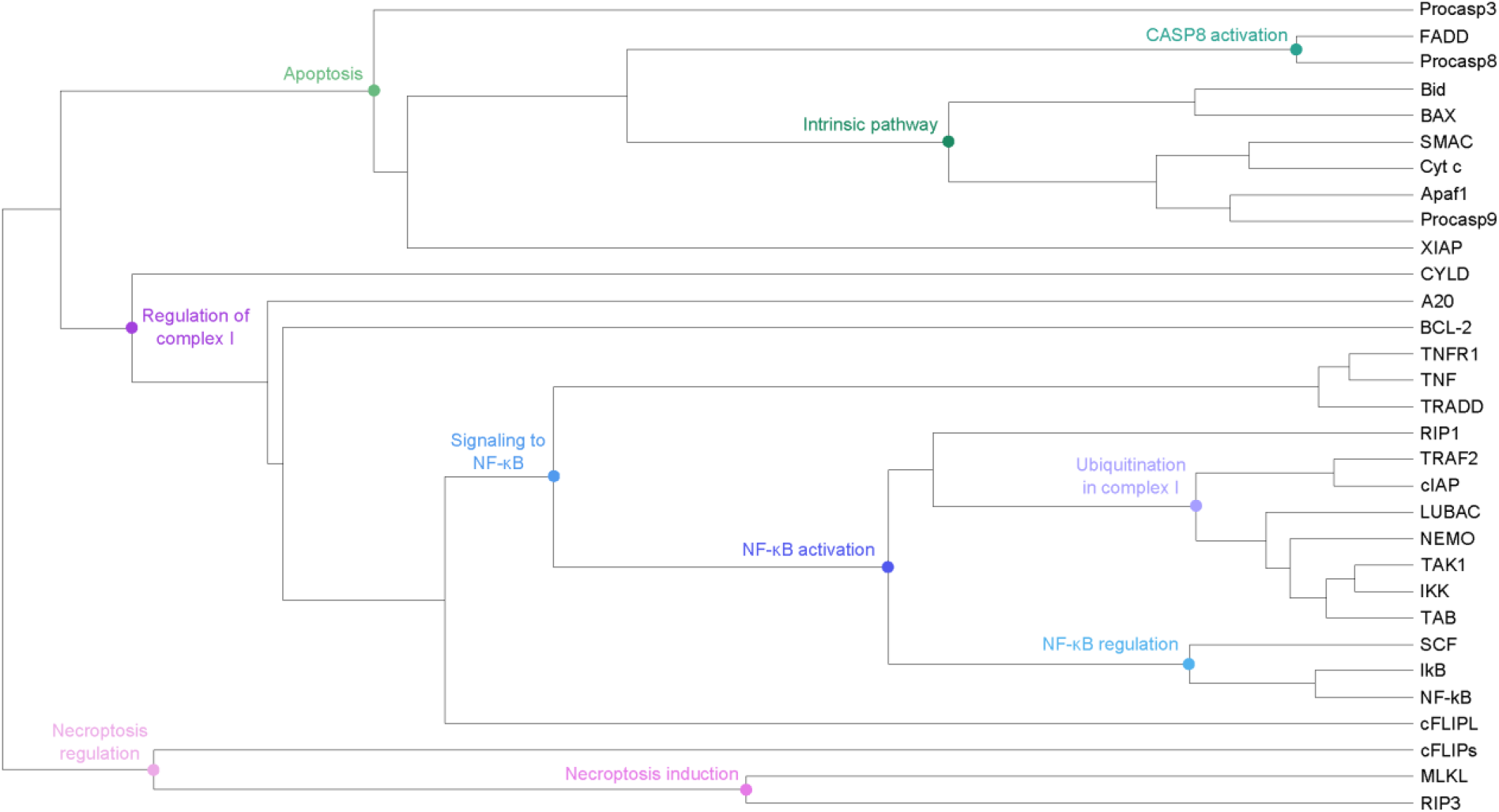
Hierarchical cluster tree. The corresponding places in the PN were clustered based on the matrix in Fig S5 in the Appendix. The hierarchical clustering was performed, using UPGMA (Unweighted Pair Group Method with Arithmetic mean) with Pearson correlation distance. Some vertices in the dendrogram were marked in blue, green, and red, referring to processes of NF-κB activation, apoptosis induction, and necroptosis induction, respectively.

### Knockout analysis of a selected submatrix

We employed the *in silico* knockout for the additional verification of the PN model of TNFR1 signal transduction in **Fig 2** and for the discussion of the molecular switch behavior. Whereas some knockout effects were obvious, others can only be derived from network analysis. **Fig 7** shows a section of the knockout matrix in **Fig S5** (Appendix). We selected twenty proteins for the single knockout and determined the effects for 21 pathway entities. The last two rows are multiple knockouts that represent the effects for Smac mimetic and the impairment of translation by cycloheximide. The selection was driven by the biological interpretation of the knockout analysis and, therefore, contains important signaling vertices in the network regarding the molecular switch. For the results, see **Table 1**.

**Table 1:** Results of *in silico* knockouts illustrated in the submatrix in Fig 7.

| Knocked out entity<br>[row no.; # red entries] | Negative effect on | Other functional aspects |
| --- | --- | --- |
| BAX [1; 6] | complex formation of BCL-2 and BAX (BCL-2:BAX); activation of CASP9 in the apoptosome (CASP9, Apoptosome) | Activation of CASP3 and CASP8 are not directly dependent on BAX (CASP3, CASP8). |
| cFLIPS [2; 1] | complex Formation of cFLIPS bound to complex IIb (CIIb:cFLIPs) | cFLIPS can promote necroptosis induction in complex IIb, but as other pathways exist that can also induce necroptosis, the knockout of cFLIPS has no direct effect on necroptosis induction. |
| cIAP1/2 and TRAF2 [3, 20; 10] | formation of complex I and the NF-κB-dependent gene expression as well as the feedback and crosstalk regulation of the target genes | This emphasizes the direct regulation of both proteins since cIAP1/2 requires TRAF2 for recruitment. |
| CYLD [4; 1] | complex formation of CYLD bound to the complex I (CI:CYLD) | CYLD promotes the dissociation of complex I and the formation of complex II. As several pathways also cover the processes, no other effects are observed. |
| FADD, procaspase 8 [5, 12; 14] | all places associated to apoptosis processes | Only the survival pathways and necroptosis induction is still functional. This emphasizes the direct regulation of both proteins since procaspase 8 requires FADD for recruitment. |
| IKK, NEMO, TAK1, LUBAC [6, 7, 9, 16; 9] | the downstream activation of NF-κB and the regulation of the target genes | Complex II formation and cell death induction remain functional. This indicates the strong relation of the proteins in the Ub-dependent regulation in complex I. |
| MLKL [8; 1] | activated MLKL located at the plasma membrane prior necroptosis induction (MLKL<sub>PM</sub>) | As activated MLKL refers to the last step in the necroptosis pathway, necroptosis induction is hampered. |
| NF-κB [10; 8] | NF-κB regulation via IκB and the regulation of NF-κB-dependent genes | |
| procaspase 3 [11; 2] | CASP3 activation and CASP3 inhibition by XIAP (CASP3, XIAP:CASP3) |  |
| procaspase 9 [13; 4] | processes of the regulation of procaspase 9 in the apoptosome via XIAP and SMAC |  |
| RIP1 [14; 14] | formation of complex I and the induction of necroptosis | Only apoptosis processes are still functioning since RIP1 is a major player in the TNFR1 signal transduction pathway. |
| RIP3 [15; 2] | formation of the necrosome and the activation of MLKL (RIP1:RIP3, MLKL_PM) |  |
| TNF, TNFR1, TRADD [17, 18, 19; 20] | all places except for the nuclear NF-κB (NF-κB_n). Since the three proteins initialize the pathway, all downstream pathway components are affected by the knockouts. | This is due to the modeling of the turnover of NF-κB, which remains unaffected by the knockout. |
| clAP1/2 and XIAP (Smac mimetic) [20; 10] | formation of complex I, NF-κB-dependent gene expression, and XIAP regulation | Only apoptosis and necroptosis induction remain functional. |
| IκB, A20, XIAP, cFLIP<sub>L</sub>, and BCL-2 (cycloheximide) [21; 7] | the translation of upregulated genes | Only the cell death pathways remain unaffected. |

**Figure 7:**
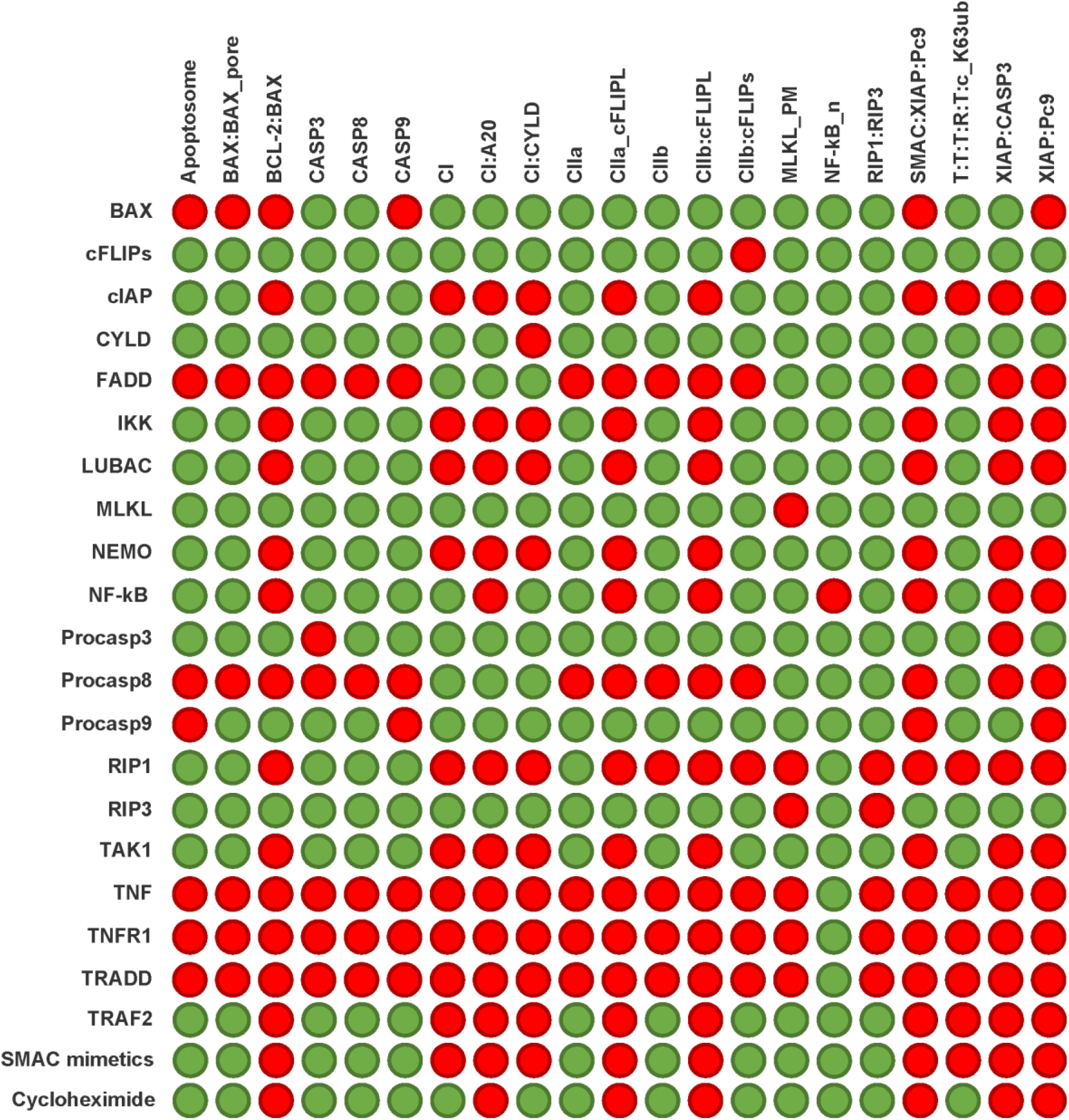
*In silico* knockout submatrix of the PN in Fig 2. The rows list the proteins, which were knocked out, and the columns give the protein complexes in the network, which might be affected by the knockout. A red (green) entry indicate that the respective complex was (was not) affected by the knockout. We performed the single knockout analysis for twenty proteins and displayed the effect for an excerpt of 21 pathway entities. The last two rows represent multiple knockouts that display the effect of Smac mimetic, i.e., the knockout of XIAP and cIAP, and the impairment of the translation of upregulated genes by cycloheximide, i.e., the knockout of IκB, A20, XIAP, cFLIP_L_, and BCL-2.

## Discussion

### The model covers signaling processes of cell survival, apoptosis, and necroptosis

The study of TNFR1 signal transduction has a long tradition and revealed many theories, depending on the current focus in the field and the respective researcher, which leads to different views and theories (Wallach, 2016; Schwabe & Luedde, 2018). The contradictions of the pathway regulation in literature and the differences of signal transduction, occurring in different cell types, require a disentangled view of the processes (Wajant & Scheurich, 2011).

The PN model compiles the current view of the TNFR1 signaling pathway, emphasizing the well-characterized processes and discussing less-known mechanisms. During the development of the model, well-established molecular regulations have been superseded by other proposed regulatory mechanisms. An example is the regulation of A20, which operates as a deubiquitinating enzyme in the feedback regulation of NF-κB signaling. Originally, its suppressive role in NF-κB signaling was assigned to the proteasomal degradation of RIP1 by a K48-linked Ub tag (Wertz & Dixit 2008). This mechanism is now questioned even though it has long been viewed as an important feedback mechanism to terminate signal transduction.

On the contrary, less-understood processes could not be integrated in the PN model as the exact mechanism of regulation is not entirely characterized. Important aspects that need further investigation are the effect of RIP1 phosphorylation and the regulation by ubiquitination within complex II. Further, the exact mechanism of necroptosis execution and the mode of action of MLKL remains to be identified.

### Model analysis reveals substance conservation, basic dynamics of the system, and all complete signaling pathways

To investigate networks of pathways in systems biology profoundly, the determination of all possible signal flows is obligatory. The mathematical approach of PIs and TIs explains substance conservation and the basic system’s behavior, repectively. TI-induced subnetworks represent functional modules. MIs constructed by liear combination of TIs represent complete functional signal flows in a network that operate at steady state. The complexity of the computation of MIs is related to the number of TIs and the possible linear combinations. For the PN of TNFR1 signal transduction, the number of MIs highly increased with regard to the number of TIs, from 48 TIs to 279 MIs.

### Knockout analysis for classification of pathways, ranking of pathway’s entities, and clustering of processes

The deduction of the regulation of signal transduction via knockout experiments is not an easy task since the pathway components are involved in several processes. Further, the variation of results between cell types, type of experiment, and working group, has an essential influence on diversity of data. The *in silico* knockout analysis can reveal obvious relations, expected dependencies, and predictions of effects that were not yet experimentally proven.

#### Pathway classification

The result is in accordance with the expected biological behavior because most cells exhibit a robust survival response and suppress the cell death induction (Ting & Bertand, 2016). The dissection of the hierarchy of a pathway is important for later use in therapeutic implication. A protein that is a player more upstream in the pathway may have also an impact on other downstream branches in an undesired form of crosstalk. Therefore, a later intervention of the pathway is often more favorable because it acts more specifically (Vandenabeele *et al*, 2010). Some proteins or complexes, which can be activated in different ways, are more robust to errors since alternative signal flows can still lead to their activation.

The components of the pathway that are involved in the processes of receptor stimulation and complex I formation, are obviously more sensitive to perturbations as many downstream branching pathways are dependent on the initialization. Therefore, TNFR1, TNF-α and TRADD are the proteins with the highest influence on other network components. Hereafter, the proteins of complex I with RIP1 are leading the way. RIP1 is an important protein, as it plays key roles in NF-κB activation, apoptosis, and necroptosis. However, not all branches of the network are RIP1-dependent, like apoptosis mediated via complex IIa. The proteins of complex I have a higher impact, too, because they have an influence on the formation of complex II, the activation of NF-κB, and subsequent gene expression. The resulting crosstalk to the cell death pathways enhances the influence of the proteins of complex I. The proteins of the intrinsic apoptotic branch, necroptosis induction as well as the proteins, which are upregulated by NF-κB, are less essential and act more specifically.

**Robustness** describes an inherent quality of systems and aims to maintain and ensure the correct function of a system (Kitano, 2004). Alternative signal flows, which target the same cellular response, enhance the robustness of a system as the function is robust to perturbations. The more redundant signal flows activate one cellular outcome, the more robust is the system to potential failing modes. The various signal flows to the different outcomes determined by MIs reveal the robustness of the TNFR1 signaling system. We concluded that the system is robust to perturbations and that the survival response is most likely to occur followed by apoptosis and then necroptosis with regard to the amount of assigned pathways.

The TNFR1 signaling pathway will always be a target of cytoprotective or cytotoxic therapies as it controls opposing responses and has a major function in immunity and development (Fulda 2011, Fulda 2014). The intertwined regulatory network makes it difficult to directly intervene cell death pathways in the desired way (Lockshin & Zekeri 2007). For cancer treatment, it is an important strategy to overcome the resistance to cell death by manipulation of signaling pathways. Such a strategy is based on Smac mimetic, which inhibits IAP proteins (Schmidt *et al*, 2018). Smac mimetic mocks the function of Smac / Diablo and inhibits cIAPs, thus, preventing RIP1 ubiquitination and phosphorylation (Ting & Bertrand 2016). It intervenes the early checkpoint and leads to a decrease of Ub chains in complex I and promotes the formation of complex II, inducing RIP1 kinase-dependent cell death (Bertrand *et al*, 2008, Fulda & Vukic, 2012). The prediction of the *in silico* knockout is in accordance with the experimental settings of Smac-mimetic treatment. Upon TNF-α stimulation, most cells do not exert cell death because of rapid gene expression of cFLIP_L_, cIAP2, XIAP, and BCL-2, which inhibit cell death signaling (Ting & Bertrand, 2016). The treatment with cycloheximide, an inhibitor of translation, or actinomycin D, an inhibitor of transcription, results in enhanced cell death (Karin & Lin, 2002).

The results of the knockout prediction may not match the experimental knockouts for every case owing to several reasons. On the one hand, the TI or MI analysis may not capture the correct pathways dependencies due to modeling reasons of abstractly modeled processes, or the knockout behavior is dependent on other signal flows, which are not explicitly included in the PN model. On the other hand, the experiments may be obtained for a specific cell type and may not be applicable to all cells. For a more predictive model with regard to the knockout behavior, we suggest to adapt the PN model of TNFR1 signal transduction to a specific cell type.

### The molecular switch

The determination of specific checkpoints of the system is important to intervene the signaling cascade in a desired manner. The survival response is very robust to perturbations as discussed above. Therefore, we needed to determine the factors that overcome this robust response and promote cell death pathways. We determined the important checkpoints in complex I in terms of the ubiquitination within complex I and the activation of NF-κB-dependent gene expression. The impairment of ubiquitination, e.g., by Smac mimetic or the *in silico* knockout of TRAF2 and cIAP, favors the induction of apoptosis and necroptois. The upregulated genes by NF-κB negatively control cell death signaling. We showed that the impairment of NF-κB activation, e.g., by knockout of proteins of complex I like LUBAC, and the translation of upregulated genes, e.g., by simulating a cycloheximide treatment, promotes cell death induction.

It is considered that ubiquitinated RIP1 has a scaffold function for the required kinases, TAK1 and IKK, in complex I and promotes cell survival (Peltzer *et al*, 2016). Deubiquitinated RIP1 can form the complex II and positively regulate cell death (Jaco *et al*, 2017; Oberst, 2017). cIAP proteins are important for TNFR1 signaling as the depletion abolishes the Ub decoration within the complex I (Tenev *et al*, 2011; Feokistova *et al*, 2011). The PN model supports this view since in absence of RIP1, only apoptosis induction can occur, and the impairment of RIP1 ubiquitination by cIAP and TRAF2 leads to the formation of complex II. Phosphorylated RIP1 is reported to inhibit kinase-dependent induction of cell death, following TNFR1 ligation (Jaco *et al*, 2017). Several studies report either IKK or MAPKAP kinase 2 (MK2), which are activated within and downstream of the complex I, to be potentially the kinases that phosphorylate RIP1 (Dondelinger *et al*, 2013; Dondelinger *et al*, 2015; Jaco *et al*, 2017; Dondelinger *et al*, 2017). It is suggested that the phosphorylation of RIP1 affects the interaction of RIP1 with FADD and CASP8 (Dondelinger *et al*, 2015; Jaco *et al*, 2017). For the association of the necrosome and the activation of RIP3, RIP1 kinase activity is required. The phosphorylation of RIP1 may function as a repressor of necroptosis besides of apoptosis (Ting & Bertrand, 2016). To integrate the exact mechanism of RIP1 phosphorylation into the PN model, further experimental studies are required.

Another checkpoint is the NF-κB-dependent gene expression, which enhances the resistance to cell death induction. Only full activation of IKK leads to NF-κB activation (Wajant & Scheurich 2011, Peltzer 2016). It was shown that the depletion or inhibition of IKK and NEMO affects the induction of apoptosis (Dondelinger *et al*, 2013; Linkermann & Green, 2014). LUBAC and TAK1 inhibition also promote complex II formation (Gerlach *et al*, 2011; Weinlich *et al*, 2016, Dondelinger *et al*, 2013). This is in accordance with the *in silico* knockout predictions for IKK, NEMO, TAK1, and LUBAC because only apoptosis induction and necroptosis induction remain functional.

The level of cFLIP_L_ is regulated by NF-κB activation. cFLIP_L_, which is a CASP8 homolog, competes with CASP8 to form a heterodimer and prevents full activation of CASP8. If NF-κB activation is blocked, the level of cFLIP_L_ decreases, leading to the induction of apoptosis (Tsuchiya *et al*, 2015). BCL-2 and XIAP are also target genes of NF-κB, which inhibit the intrinsic apoptosis pathway and apoptosis induction by caspase inhibition, respectively (Shore & Nguyen, 2008). Cycloheximide treatment impairs the translation of upregulated genes and the knockout, which represents the treatment, is in accordance with the expected behavior.

Whereas the checkpoints that mediate signal transduction in complex I and from complex I to complex II are quite well-characterized. The exact regulation within complex II is not entirely clarified. In complex II, the checkpoints mainly control caspase activity. TRADD needs to dissociate from complex I and binds to FADD to provide a platform for CASP8 recruitment and apoptosis induction (Micheau & Tschopp 2003). cFLIP_L_ is usually upregulated by the time that complex II can form in the cytosol and inhibits caspase activation. The two isoforms of cFLIP differentially regulate the activity of complex II (Feokistova *et al*, 2011). While cFLIP_L_, binding to CASP8 and FADD has a survival function, blocking apoptosis and necroptosis, cFLIPS binding to CASP8 inhibits full activation of caspase activity (Oberst *et al*, 2011; Dillon *et al*, 2012). There are evidences that the formation of complex IIa and complex IIb has also several checkpoints involving post-translational modifications. The influence of ubiquitination in complex II needs to be further studied (Onizawa *et al*, 2015). CYLD is a substrate of CASP8, which may be involved in the regulation of the switch of complex IIa to complex IIb (O’Donnell *et al*, 2011). Also A20 is reported to inhibit RIP3 activation by ubiquitination and prevents necroptosis induction, which would result in another crosstalk from the target gene of NF-κB (Onizawa *et al*, 2015).

## Materials and Methods

### Petri nets

Petri nets (PNs) represent a graph theory-based mathematical formalism to model systems of concurrent processes (Reisig, 1985; Murata, 1989). PNs are widely used in technical applications (for a review, see Zhou & Azlan, 2016) and systems biology (for example, Reddy *et al*, 1993; Koch *et al*, 2005; Sackmann *et al*, 2007; Grunwald *et al*, 2008; Kielbassa *et al*, 2009; Koch *et al*, 2011; Koch *et al*, 2017). PNs are directed, bipartite, labeled graphs. There are two type of vertices, one type for the passive elements of the system called places and one for the active elements called transitions. For biochemical systems, the places model biological entities, for example, proteins, RNAs, ligands, protein complexes, genes, and other chemical compounds. The transitions stand for the reactions transforming one place into another, for example, chemical reactions, phosphorylation, ubiquitination, complex formation, and other. The directed edges connect only vertices of different type. Places with outgoing edges are called pre-places and places with ingoing edges post-places, with respect to the transition the edges connect with the condidered place. Edges can be labeled, usually by integers.

Formally, we define a PN as a quintuple N = (*P, T, F, W, m*_0_) with:

*P* = {*p*_1_, *p*_2_,…, *p*_*m*_} is the finite set of places.

*T* = {*t*_1_, *t*_2_,…, *t*_*n*_} is the finite set of transitions.

*F* ⊆ (*P* × *T*) ∪ (*T* × *P*) is the set of flow relations or edges.

*W* : *F* → N defines the edge weights.

*m*_0_ : *P* → N_0_ is the initial marking.

For classical PNs called P/T nets (Place/Transition nets), the dynamic is performed by movable objects named tokens that are located on the places. A token represents a discrete entity, for example, one mole of a chemical compound or one molecule. A certain token distribution defines a specific system state and is given by the marking, *m*, which is a vector of the size of *P*. The initial marking, *m*_0_, describes the initial state of the system before starting a simulation. The marking is illustrated by points on the corresponding places or by the numbers.

Tokens move through a PN following specific friring rules. In P/T nets. These firing rules are timeless, meaning that the tokens on the pre-places are removed at the same time as the tokens are produced on the post-places, see **Fig 8**.

**Figure 8:**
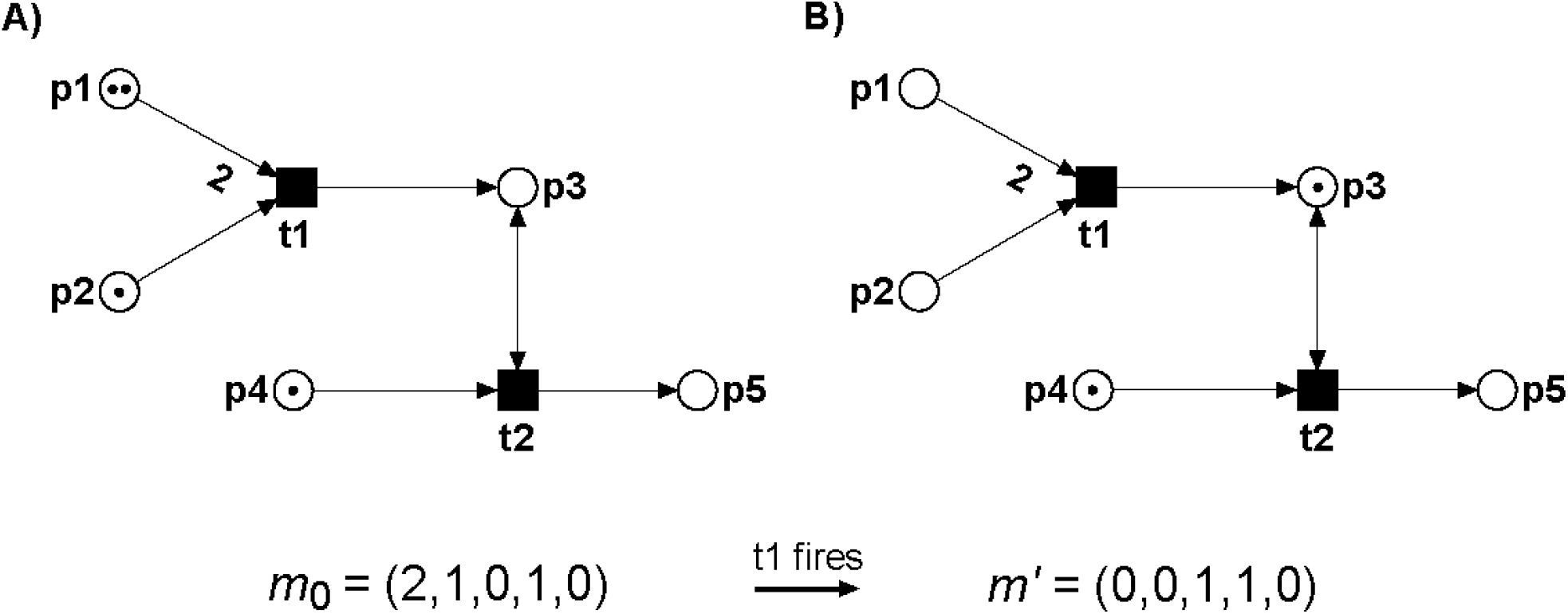
PN example 1. A) The PN consists of five places depicted by circles, two transitions depicted by rectangles, and six directed edges. The edge from p1 to t1 has a weight of two. For all other edges with a weight equals one, no label is drawn. p3 and t2 are connected via a read arc. Tokens are depicted as dots on the places p1, p2, and p4, defining the initial marking *m*_0_ = (2, 1, 0, 1, 0). B) The PN after the firing of t1. The marking has changed by removing tokens from the pre-places, p1 and p2, and producing a token on the post-place, p3. Then, the new marking is *m*’ = (0, 0, 1, 1, 0).

We modeled the PN as an open system, meaning that all proteins of the pathway are synthesized and degraded. The only exceptions are the genes that induce the synthesis of proteins in a controlled manner and, therefore, form specific patterns in the PN model.

### Invariants

Among other properties, a PN is characterized by its invariants – a property that always holds at steady state independent of the system state and the initial marking. Invariants can relate the structure of the net to the behavior of the system and allow for an implication of the system’s dynamics.

We define transition invariants (TIs) and place invariants (PIs). Both are based on the incidence matrix of the PN, *C* = *P* × *T*. An entry in *C* indicates the change in the number of tokens on the considered place (row), if the considered transition (column) fires. Based on the incidence matrix, we can define two invariant properties, transition invariants and place invariants.

### Transition invariants

A TI is a set of transitions, whose firing sequence reestablishes an arbitrary initial marking, Δ*m*_0_ = 0, i.e., the system state is invariant. A TI is defined as a Parikh vector, *x*, that fulfills the equation Δ*m* = *C x* = 0. The exact number of firings per transition is given by the elements of *x*. An integer solution is a true invariant, *x*, if x has no negative components, i.e., *x* ≥ 0. The set of transitions, whose components in *x* are positive, defines the support of the TI, supp(*x*). A TI is minimal if no other solution, *x*’, exists with supp(*x*’) ⊇ supp(*x*), and the largest common divisor of all elements equals one. A PN is *covered by TIs* (CTI) if every transition is member of at least one TI.

### Place invariants

Analogously to TIs, we define PIs of a PN as a vector, *y*, applying *C*^T^ *y* = 0, where *C*^T^ denotes the transposed incidence matrix. The definition of a minimal and true PI is analogous to the definition of a minimal and true TI. A PN is *covered by PIs* (CPI) if every place is member of at least one PI.

### Manatee invariants

MIs are linear combinations of TIs to ensure that the TI covers a signaling pathway from receptor activation to cell response. For the detailed definition of MI, we refer to Amstein *et al*, 2017.

### Invariant-induced subnetworks

Each invariant induces a subnetwork. A TI-induced or MI-induced subnetwork is formed by the transitions of the TI or MI, respectively, and the places and edges in between. Analogously, a PI-induced subnetwork is defined by the places oft the PI and transitions and edges in between.

### *In silico* knockout analysis

Knockout studies or perturbation studies are suitable methods to reveal the vulnerable parts of a model. The *in silico* knockout analysis supports a profound investigation of a comprehensive PN model of a signaling pathway (Scheidel *et al*, 2017). We define a knockout matrix, where each row represents the knockout of a protein, i.e., the deletion of an input transition. Each column indicates proteins or protein complexes of the PN, which might be affected by the considered knockouts. We visualize the knockout results by coloring the matrix entries either green (red) if the place is part (not part) of at least one MI-induced subnet. Biologically, the green entry indicates that the respective protein or protein complex remains unaffected by the knockout, while a red entry stands for an effect on protein or protein complex formation.

We ranked the proteins of the TNFR1 pathway according to their influence on other pathway components based on the knockout analysis performed for the complete set of proteins and protein complexes. We selected all transitions, which represent protein syntheses, and all places of the PN model except for the places belonging to a PI. The knockout was performed applying isiKnock (Hannig *et al*, 2019) based on MIs using additional output transitions.

Moreover, we performed a clustering of the knockout data of the complete knockout matrix in **Fig S5** in the Appendix. For the hierarchical clustering of the matrix entries, we applied the software NOVA with the settings UPGMA with Pearson correlation distance (Giese *et al*, 2015).

## Acknowledgements

This work was partly supported by the Cluster of Excellence ‘Macromolecular Complexes’ of the DFG (3212070002/TP2) and by the LOEWE program Ubiquitin Networks (Ub-Net) of the State of Hesse (Germany) (20120712/B4). The founders had no role in study design, data collection and analysis, decision to publish, or preparation of the manuscript. We acknowledge and thank the Goethe-University Frankfurt am Main and Hessian Ministry of Higher Education, Research and the Arts for providing financial and infrastructural support.

## Author contributions

ID, IK, and SF conceived the study. JA, LA, and IK designed the computational analyses. LA and JH collected the data. LA, JH, and JA developed the Petri net models and performed the analyses. ID, IK, and SF provided funding and supervised the study. LA, IK, and JA wrote the paper. All authors approved the publishing of the manuscript.

## Conflict of interest

The authors declare that they have no conflict of interest.

